# Randomized controlled studies comparing traditional lectures versus online modules

**DOI:** 10.1101/2021.01.18.427113

**Authors:** Kiran Musunuru, Zarin P. Machanda, Lyon Qiao, William J. Anderson

## Abstract

We assessed the efficacy of traditional lectures versus online modules with respect to student learning in an undergraduate introductory biochemistry course in two successive years. In the first year, students had the options of attending live lectures by the course instructor and viewing online modules pre-recorded by the instructor, with the lectures and modules covering identical content; in addition, all students had a mandatory weekly application session. Utilizing pre-course and post-course tests as an instrument with which to measure learning during the course, we observed significantly increased learning (0.7 standard deviations) with attendance of traditional lectures and decreased learning with use of online modules, even after adjustment for grade point average. In the second year, the course had the same curriculum, but students were randomized to either live lectures or online modules for the first half of the course, crossing over to the other modality during the second half. With randomization, no difference in learning was observed between the two groups. Furthermore, we found that students self-reported greater engagement when viewing online modules than when attending lectures in person. These findings suggest some aspects of the lecture experience can be shifted to online modules in STEM courses without impacts on student learning so as to use classroom time more fully for application-based active learning interventions.

## INTRODUCTION

Active learning interventions have been persuasively demonstrated to improve student performance in STEM courses (Mazur, 1997; Wood, 2003; Knight and Wood, 2005; Smith et al., 2009; Wood, 2009; Smith et al., 2011; National Research Council, 2012; Freeman et al., 2014). Active learning is defined by Driessen et al. (2020) as “*an interactive and engaging process for students that may be implemented through the employment of strategies that involve metacognition, discussion, group work, formative assessment, practicing core competencies, live-action visuals, conceptual class design, worksheets, and/or games*”. As such, it has been advocated that STEM education research should henceforth focus on comparing the efficacy of different active learning interventions rather than using traditional lectures as the comparison standard, with lectures likened to the “pedagogical equivalent of bloodletting” (Freeman et al., 2014; Wieman, 2014). The increasingly popular flipped classroom model, in which active learning sessions are paired with online modules that students are expected to have viewed before the sessions, allows for classroom time to be devoted entirely to active learning approaches. Perforce, the flipped classroom model assumes that traditional live lectures by a “sage on the stage” have less value for student learning, and that introduction of course material occurs just as readily, if not more readily, via online instruction (King, 1993; Lage et al., 2000; Parslow, 2012).

Some notable examples from the literature drew our attention, starting with an early meta-analysis of visual-based instruction describing a slight advantage of videos versus traditional lectures in the pre-Internet era (Cohen et al., 1981). Student performance in an introductory human-computer interaction course was significantly higher in a cohort that solely relied on online lectures as opposed to in-class lectures using both grades and student attitudes as readouts (Day and Foley, 2006). Similar results were seen in an introductory biology course that employed pre-recorded videos or pre-class worksheets to replace components of a subset of lectures (Moravec et al., 2010). We also came across reports using randomized controlled studies comparing online versus live lectures (Porter et al., 2014; Vaccani et al., 2016; Brockfeld et al., 2018; Chirikov et al., 2020). While the student populations differed in terms of level (undergraduate versus graduate), discipline (engineering, pharmacy, medicine), and duration of the course (one week to a full semester), the results were similar, showing no difference in learning outcomes with online lectures compared to synchronous in person lectures.

We sought to assess whether the same trend held true in a semester-long introductory biochemistry course that covered the topics now included in the Medical College Admissions Test, which was reformulated in light of recommendations from the American Association of Medical Colleges and the Howard Hughes Medical Institute (AAMC-HHMI, 2009; Alpern et al., 2011). During the first year the course was taught, we offered students the choices of whether to attend in-class lectures and whether to view online modules covering the same material. In the second year, we switched to a different study design in which students were randomized to in-class lectures or online modules at the beginning of the semester and then switched halfway through the semester, allowing us to embed two randomized controlled studies within the semester-long course. In both years, pre-course and post-course tests were used to measure learning gains. The contrasting study designs of the two years allowed us to explore not only the relative efficacies of the two learning modalities but also whether student motivation to be in the classroom influences their degree of learning.

## RESULTS

### Course Design

Study activities described herein occurred at Harvard University in Cambridge, Massachusetts. The course, entitled *SCRB 25: Biochemistry and Human Metabolism* and taught during two consecutive spring semesters, combined expository instruction and its application. Lectures comprised 1.5-hours per week, with all of the lectures given by the course instructor (the first author of this manuscript) early in the week. The course instructor also narrated online modules covering the identical material, showing the lecture slides in the format used by the Khan Academy (i.e., only the slides are seen in the video, not the instructor) (Parslow, 2012). The online modules were made available each week on the course website shortly before the live lectures. As much as possible, the instructor attempted to standardize his delivery style in the lectures and the online modules. In general, each week’s material was split into two to three online modules, mirroring the lectures being split into blocks with short intervening breaks. When students asked questions during lecture, the instructor committed to making a transcript of the questions and their answers available to all students enrolled in the course.

Because we wished for all students to benefit from synchronous active learning interventions with their peers, the course also had a mandatory, weekly 1.5-hour section (ranging from 16 to 20 students in each section during the first year, and from 27 to 32 students in each section during the second year) later in the week in which students were randomly split into small peer groups (three to four individuals in each group) to work through patient cases intended to illuminate the various principles of biochemistry covered in the lecture/online modules earlier that week (see Supplement for an example patient case). Teaching assistants provided guidance for the peer groups, while the groups discussed questions provided with the cases, and facilitated section-wide discussion of the answers to the questions. The sections additionally provided the opportunity for students to ask the teaching assistants about course material. The teaching assistants as well as the course instructor also held regular office hours each week for this purpose. Students were evaluated with problem sets, assigned on average every two weeks, and midterm examinations. The course enrollment was 86 students during the first year and 145 students during the second year, in each case a mix of sophomores, juniors, and seniors.

### Uncontrolled Study in the First Year

Our goal with the course design was to compare the efficacy of traditional lectures versus online modules with respect to student learning. We asked students to choose between the two modalities, allowing for the possibility of individual students attending the lectures during some weeks and viewing the online modules during other weeks, or even doing both (or neither).

We developed a 20-question test that was representative of the entire semester’s course material. The questions were adapted from questions provided by W. H. Freeman and Company, the publisher of *Lehninger Principles of Biochemistry*, Sixth Edition, the textbook used for the course. We administered the same test during the active learning sections in the second week of the semester (the first meeting of the sections) and during the last week of the semester. Students were given 20 minutes to complete the set of 20 questions in closed-book testing conditions. It was made clear to the students that the pre- and post-course tests were optional, had no bearing on the course grade, and were being administered for the purpose of a research study. A total of 72 students completed both the pre-course test and the post-course test.

In addition to the 20 biochemistry questions, students were asked additional information: college year, whether they had taken or were concurrently taking another biochemistry-related course, to rate on a scale of 1 to 10 how often they had attended live lectures during the semester (Fig. 1B), and to rate on a scale of 1 to 10 how often they had viewed the online modules during the semester (Fig. 1C). Separately from the tests, students filled out surveys to express their thoughts about the course. Subsequent to the end of the semester, we obtained the students’ cumulative grade point averages (GPAs) from the university registrar.

**Figure 1.**
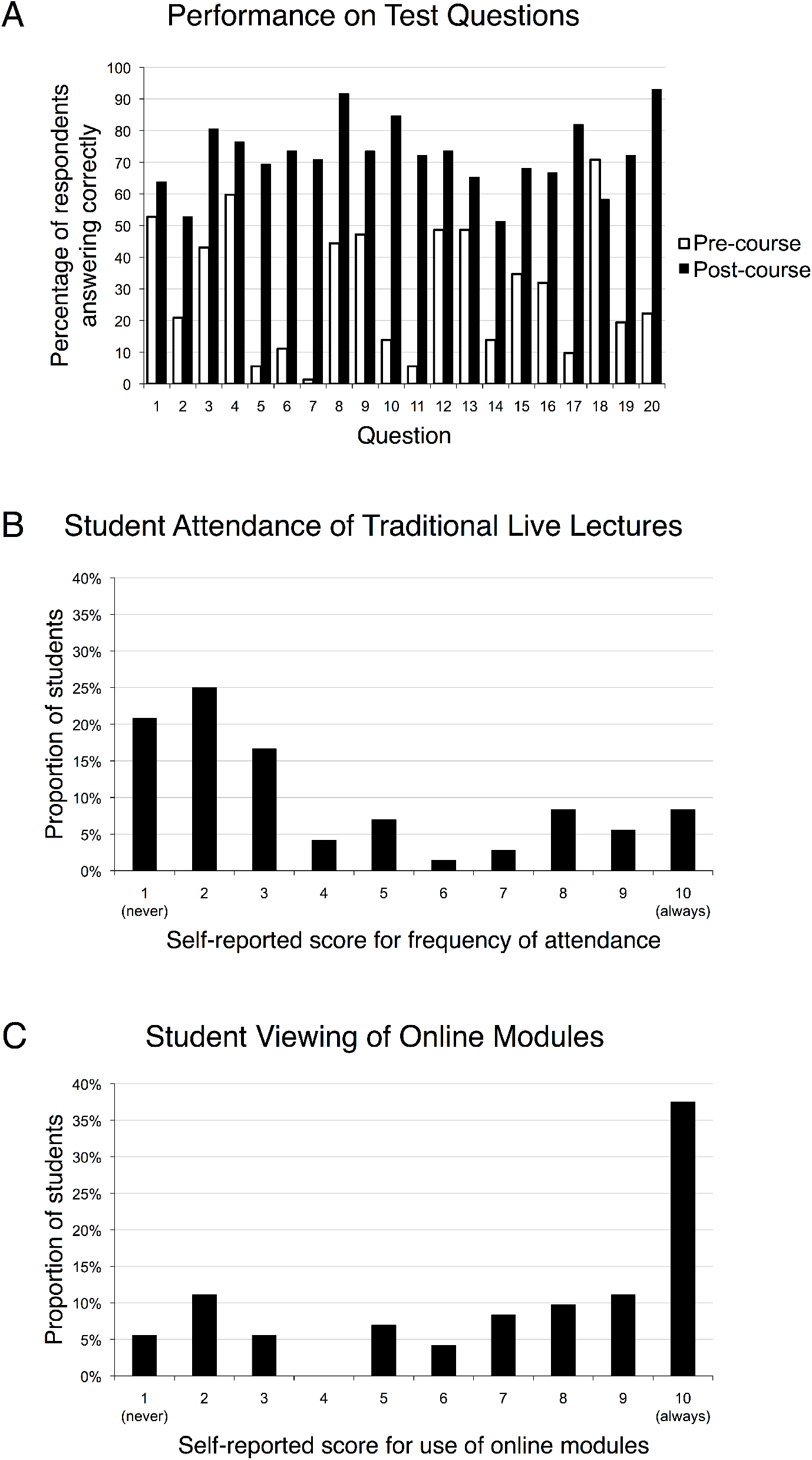
Test performance and attendance of lectures versus use of online modules during the first year. (**A**) Percentages of correct answers for each of the 20 questions on the pre-course/post-course test. (**B**) Proportions of students who rated their attendances of lectures on a scale of 1 (none) to 10 (all). (**C**) Proportions of students who rated their use of online modules on a scale of 1 (none) to 10 (all).

Utilizing the pre-course and post-course tests as an instrument with which to measure learning during the course, we observed an increase in scores from a mean of 30% to a mean of 72%. The percentage of respondents correctly answering each question improved for 19 out of the 20 questions (Fig. 1A). We calculated the normalized learning gain—the difference between the post-course test score and pre-course test score divided by 100% minus the pre-course test course—to represent the raw gain divided by the maximum possible gain for each student, thus taking into account the variation in pre-course knowledge of biochemistry (Wood, 2009). Every single student experienced a positive learning gain.

We found that more students chose to rely exclusively or predominantly on the online modules than on live lectures (Fig. 1B and 1C), with a few students attending some lectures and viewing some online modules. As expected, there was a strong inverse correlation between attendance of lectures and use of online modules (Pearson’s *r* = –0.47, *P* = 0.00004). Using simple linear regression models (Table 1), we found a significant positive relationship between the degree of attendance of lectures and learning gain (*P* = 0.04). Based on the models, we found that attendance of all lectures resulted in an absolute 16% increase in learning gain (71% vs. 54%). There was a similar change in learning gain associated with the use of online modules, but in the opposite direction (55% vs. 71%). The standard deviations (SDs) of the learning gains for all participants was 23%; thus, the absolute increase in normalized learning gain observed with attendance of lectures corresponded to an effect size of 0.7-SD.

**Table 1.** Associations of variables with learning gains in the first year.

| Variable | Normalized learning gain<br>(*P* value) | Effect<br>(regression coefficient) |
| --- | --- | --- |
| Attendance of lectures* | 0.04 | +1.83 |
| Use of online modules* | 0.04 | −1.77 |
| Cumulative grade point average<sup>†</sup> | $7 \times 10^{-6}$ | +47.73 |
| College year | 0.25 |  |
| Previous enrollment in biochemistry-related course | 0.29 |  |
| Concurrent enrollment in biochemistry-related course | 0.02 |  |
| Attendance of lectures*<sup>‡</sup> | 0.02 | +1.68 |

\*Rated on a scale of 1 (none) to 10 (all)
<sup>†</sup>Calculated on a four-point scale (0.00 to 4.00)
<sup>‡</sup>Adjusted for cumulative grade point average and concurrent enrollment in biochemistry-related course

We tested several other variables including college year, previous enrollment in a biochemistry-related course, concurrent enrollment in a biochemistry-related course, and cumulative GPA. We found that only GPA and concurrent enrollment showed evidence of association with normalized learning gain (Table 1). While GPAs were strongly correlated with learning gains, we did not observe significant correlation of GPA with attendance of lectures (Pearson’s *r* = 0.10, *P* = 0.41) or with use of online modules (Pearson’s *r* = –0.11, *P* = 0.35). In generalized linear models that included attendance of lectures, GPA, and concurrent enrollment, attendance of lectures remained significantly associated with the learning gain (*P* = 0.02), with the effect size essentially unchanged.

### Randomized Controlled Studies in the Second Year

In the second year the course was offered, we used exactly the same course materials and format of lectures/online modules paired with application-based sessions, but we switched to a controlled study design in which students were randomized to in-class lectures or online modules. To encourage students to participate in the study, we adopted a crossover design so that all participants had the same cumulative exposure to in-class lectures and online modules. Additionally, the cumulative final exam for the course was optional for the participants. A total of 124 out of 145 students (86%) participated in the study.

Participants were randomized at the end of the first week of the course and took a pre-test of 20 questions covering the material for the first half of the course. The test included questions used in the test from the first year of the course (the questions relevant to the material covered in the first half of the course) with the addition of new questions related to the material. For weeks two through six, the students randomized to the in-class lectures were encouraged to come to class, where attendance was taken solely for the purpose of the study (i.e., was not factored into the student grade) and did not have personal access to the online modules for those weeks on the course’s learning management system. During the same timeframe, the students randomized to the online modules had free access to the modules on the learning management system but were not permitted to attend the in-class lectures. Near the midpoint of the course, in the week prior to the first midterm examination, the students took a combined post-test for the first half of the course and pre-test of 20 questions for the second half of the course (as with the test for the first of the half of the semester, the test for the second half was a mix of old questions and new questions). At that point, the participants crossed over to the other learning modality, with the same restrictions on lecture attendance or online module access during weeks seven through twelve. Near the end of the course, in the week prior to the second midterm examination, the students took a combined post-test for the first half of the course (to assess for durability of learning) and post-test of 20 questions for the second half of the course.

Because we had effectively performed two randomized controlled studies during the semester, we had four measures available with which to compare the two learning modalities: (1) the learning gain for the first half of the course, as judged by the pre- and post-test scores; (2) the grade on the midterm examination for the first half of the course; (3) the learning gain for the second half of the course; and (4) the grade on the midterm examination for the second half of the course. On all four measures, we did not observe any significant differences between the in-class lecture group and the online module group (Table 2). We also found that the measured learning gains for the entire class during the second year of the course were less than the measured learning gains during the first year of the course (Table 2). This may reflect that the learning period was a full semester in the study performed in the first year, whereas the learning periods were only half-semesters in the study performed in the second year. We did find that the learning gains during the first half of the semester in the second year were retained through the end of the semester and, indeed, were slightly larger, perhaps reflecting that the material covered during the second half of the semester (more focused on metabolism) helped reinforce the learning that occurred during the first half (more focused on biochemistry).

**Table 2.** Learning gains and midterm grades in randomized controlled studies.

|  | Normalized learning gain (median) |  |  | Midterm grade (median)* |  |  |
| --- | --- | --- | --- | --- | --- | --- |
|  | Traditional lectures | Online modules | *P* value<sup>†</sup> | Traditional lectures | Online modules | *P* value<sup>†</sup> |
| Pre-test vs. post-test 1 (semester midpoint) for 1st half of semester | 0.32 | 0.35 | 0.98 |  |  |  |
| Pre-test vs. post-test 2 (semester end) for 1st half of semester | 0.38 | 0.37 | 0.38 |  |  |  |
| Pre-test vs. post-test for 2nd half of semester | 0.35 | 0.29 | 0.54 |  |  |  |
| Midterm for 1st half of semester |  |  |  | 43.5 | 44.0 | 0.70 |
| Midterm for 2nd half of semester |  |  |  | 46.5 | 46.5 | 0.70 |

\*Scored out of 50 points
<sup>†</sup>Calculated with Mann-Whitney *U* test

### Additional Information Obtained from the Studies

In student surveys from the first year, when asked to express a preference for traditional lectures or online modules, only 32% chose traditional lectures, with 68% choosing online modules— consistent with the actual behavior of the students when offered the choice during the first year. In student surveys from the second year—administered after the completion of the randomized controlled studies—when asked to express a preference for traditional lectures or online modules, 40% chose traditional lectures, with 60% choosing online modules; when asked which modality they believed was best for their learning, 23% chose traditional lectures, 54% chose online modules, and 23% thought that both were equally effective. Interestingly, in student surveys from the second year, 52% indicated that they would have both attended in-person lectures and used the online modules if both modalities had been available to them. This is at odds with actual student behavior during the first year of the course, when the students did have both modalities available to them—very few students took advantage of both.

We attempted to ensure that students adhered to their assignment group. Attendance at lectures was carefully recorded by teaching assistants, and on no occasion was a student assigned to the online modules present at a lecture. However, among the students assigned to the lecture group, a substantial number of them missed lectures; averaged throughout the semester, about 60% of the students assigned to the lecture groups were in attendance at any given lecture. Although we attempted to restrict viewing of the online modules to just those students assigned to the modules via personal logins in the learning management system, 22% of the students reported that they were nevertheless able to find ways to watch modules when assigned to the lecture group, presumably by having classmates assigned to the online module group give them access to the modules. These factors would have weakened the power of the randomized controlled studies to detect a difference in learning from traditional lectures versus online modules.

Another important consideration is the extent of engagement of the students when in attendance at lectures or while viewing online modules. In student surveys from the second year, students self-reported the following activities during lectures, at least to an occasional degree: 87% checked email; 71% either chatted online or texted with their mobile devices; 65% surfed the Web; 59% talked with classmates; and 35% worked on other assignments. Students self-reported the following activities during the viewing of online modules, at least to an occasional degree: 66% checked email; 57% surfed the Web; 56% either chatted online or texted with their mobile devices; 27% talked with others in person or by telephone; and 19% worked on other assignments. We asked students to quantify each of the aforementioned activities on a scale of 1 (never), 2 (rarely), 3 (sometimes), 4 (often), and 5 (always). We found that for all of the five activities, the students engaged in each activity to a lesser degree when watching online modules than when attending lecture, with three of the activities—checking email, talking with others, and working on other assignments—having statistically significant differences (Table 3). Of note, 77% of students reported taking advantage of the ability of the learning management system to play the online modules at 1.5× or 2× speed, potentially reducing the amount of the time spent in the learning activity.

**Table 3.** Activities while attending traditional lectures or viewing online modules.

|  | Traditional<br>lectures* | Online<br>modules* | *P* value<sup>†</sup> |
| --- | --- | --- | --- |
| Checked email | 2.86 ± 1.14 | 2.23 ± 1.12 | 1 × 10<sup>-5</sup> |
| Chatted online or texted | 2.37 ± 1.19 | 2.26 ± 1.15 | 0.58 |
| Surfed the Web | 2.24 ± 1.20 | 2.07 ± 1.13 | 0.21 |
| Talked with others | 2.03 ± 1.10 | 1.47 ± 0.85 | 2 × 10<sup>-5</sup> |
| Worked on other assignments | 1.60 ± 0.98 | 1.28 ± 0.66 | 0.003 |

\*Mean and standard deviation; scored on a scale of 1 (never), 2 (rarely), 3 (sometimes), 4 (often), and 5 (always)
<sup>†</sup>Calculated with Wilcoxon signed-rank test

## DISCUSSION

In the uncontrolled study we performed in the first year of the course, we observed increases in learning gains with attendance of lectures corresponding to an effect size of 0.7-SD. This compares favorably with the average 0.47-SD improvement reported for active learning interventions in STEM courses (Freeman et al., 2014). We considered two possible explanations for the increased student learning seen with increased attendance of lectures (or, conversely, the decreased student learning seen with increased use of online modules). First, the in-class lectures resulted in superior learning compared to online modules. Second, the students who chose to attend the lectures were intrinsically more motivated to learn, i.e., a form of self-selection bias. We sought to address the latter possibility by including student GPAs in the analyses. However, we found no correlation between GPA and attendance of lectures, and the strong association between attendance of lectures and student learning endured after adjustment for GPA.

To distinguish between the possible explanations, we undertook a randomized controlled study design during the second year of the course, which would either confirm or refute that the in-class lectures resulted in superior learning compared to the online modules. The randomized controlled studies in the second year were as closely matched as feasible to the uncontrolled study in the first year, in the sense that the course material was exactly the same (and taught by the same instructor) and similar testing instruments were used. Contrasting with the uncontrolled study in the first year, in the randomized controlled studies no differences were observed in student learning between the group attending lectures and the group using the online modules. This is similar to other randomized studies that examined performance related to material in a given lecture(s) delivered in person versus online (e.g., Spickard et al., 2002; Solomon et al., 2004), as well as randomized controlled studies that looked at courses up to a semester long (Porter et al., 2014; Vaccani et al., 2016; Brockfeld et al., 2018; Chirikov et al., 2020).

One surprising finding of the study in the second year related to the degree of student engagement with the two learning modalities. Because the randomized controlled study design mandated that all participating students spend time engaged in both attending in-class lectures and viewing online modules, each student was able to make an informed self-assessment as to how often s/he engaged in “distracting” activities with both learning modalities. We found that, across the board, students spent less time on such activities when viewing online modules than when attending lectures. While we caution that we did not attempt to validate student self-report on their behaviors, this would suggest that when given the flexibility of choosing when to engage with the material, students may pick times that allow them to be more focused.

There are limitations to our study. First, in terms of the course setup, the lecture component was didactic with little active learning components, in stark contrast to the mandatory application sections attended by all students. Thus, it is possible that our results may have differed if active learning interventions were a predominant part of both the in-person and online lecture modules. Second, we used GPA as a measure of student motivation. This is not an ideal measurement, as there are specific tools designed to measure student motivation (see for example Pintrich and DeGroot, 1990; Pintrich et al., 1991; Glynn et al., 2011). Also, there are numerous reports demonstrating how active learning closes the achievement gap between students (see for example Beichner et al., 2007; Theobold et al., 2020). Third, we relied on student self-report for scoring student behavior in terms of assignment group adherence and frequency of distracting activities. These self-reports were not validated independently (e.g., through the use of software) and so they may be inaccurate. Fourth, the statistical analyses presented herein could have greatly benefitted from an increased sample size. Finally, these data may be misinterpreted to mean that the in-person classroom experience can be mimicked in its entirety online. We wish to emphasize that in this study, all students, regardless of their assigned group, met in-person for case-based collaborative learning activities in which they applied material from lecture and were able to work with their peers with teaching staff present to facilitate.

The past decade or so has seen the emergence of massive open online courses (MOOCs). Although the main benefit of MOOCs is their making college-level education available to large numbers of people for whom it was not previously accessible, MOOCs have also provided an impetus for colleges to adopt the flipped classroom model by “blending” online modules (whether custom-made or adapted from MOOCs) with active learning sessions. Prior studies suggest that online videos may also help to reduce the achievement gap between low and high performing students (Murphy and Stewart, 2015). Our results, along with others, suggest that the replacement of parts of traditional lectures with online modules in STEM courses would not compromise student learning and may possibly promote student engagement with the course material.

Importantly, given the shift to online instruction brought about by the COVID-19 pandemic, some students and faculty alike may have felt concerned as to whether student learning would be negatively impacted, especially for those students who, for various reasons (e.g., time zone differences, work/family obligations, etc.), may not be able to participate in lectures synchronously. Our study, as well as others, suggest that student learning can be similar if parts of a course are asynchronous in nature and active learning is a central tenant.

We do not believe, nor advocate, that in-person, synchronous instruction is dispensable for optimal student learning. Rather, we would argue that asynchronous online aspects may be successfully incorporated into a course without serious impact on student learning if designed correctly, so that more active learning activities can be part of the classroom experience. We believe, as has been unambiguously and consistently demonstrated in the literature, that synchronous active learning interventions that allow students to interact with their peers and engage more deeply with the material is a hallmark of learning in the sciences. There are currently many ways in which active learning interventions can be a part of online content, even if delivered asynchronously. For example, having short online modules that are broken up with clicker-style questions allow students to test their understanding of material (see Smith and Knight, 2020 for guidelines on writing effective questions). Software like Panopto allows one to easily add these questions into recorded videos. Others have reported ways to engage students in online problem solving (see for example Anderson et al., 2008). For synchronous online lectures, utilizing breakout rooms in Zoom allow for think-pair-share activities and to work in groups using case-based collaborative learning. The COVID-19 pandemic is an opportunity to utilize these online tools for the betterment of our courses so that online content can aid our in-person teaching when we return in person to the classroom.

## METHODS

The research described in this study was approved by the Harvard Committee on the Use of Human Subjects (protocols #IRB14-0155 and #IRB14-4563). Statistical analyses were performed using SPSS v. 21. In the uncontrolled study in the first year, simple linear regression models were used to test each variable against the normalized learning gains. Generalized linear models were used to incorporate attendance of lectures, concurrent enrollment in a biochemistry-related course, and cumulative grade point average to explain the normalized learning gains. Attendance of lectures, use of online modules, and cumulative grade point average were treated as continuous variables. College year, previous enrollment in a biochemistry-related course, and concurrent enrollment in a biochemistry-related course were treated as categorical variables. Courses considered to be related to biochemistry were *Chemistry 27: Organic Chemistry of Life*; *MCB 56: Physical Biochemistry: Understanding Macromolecular Machines*; *MCB 234: Cellular Metabolism and Human Disease*; and *BIOS S-10: Introduction to Biochemistry* (summer school course). *P* values for the variables in linear regression models were calculated with *t*-tests except for college year, for which ANOVA was used.

For the randomized controlled studies in the second year, students were randomized by assigning a random number using the random sequence generator on Random.Org (https://www.random.org/sequences) to each student name listed alphabetically. Students were then sorted by ascending value of their random number. The first half of the students were assigned to lecture only for the first half of the term, while the second half were assigned to online modules only for the first part of the term.

The Mann-Whitney *U* test was used to compare learning gains and midterm scores between the randomized groups. The Wilcoxon signed-rank test was used to compare the extent to which students engaged in activities during the in-class lectures and online modules.

## ACKNOWLEDGMENTS

We are most appreciative of the kind feedback from prior reviewers who contributed thoughtful advice and criticism of this work. We would like to thank the students of *SCRB 25: Biochemistry and Human Metabolism* for their willingness to participate in the studies; the teaching assistants for their help in collecting data for the studies. We thank our colleagues at Harvard for their support in helping to execute the study, including the Harvard Committee on Undergraduate Educational Policy; Genevieve Saphier and Kristen Elwell for their assistance with compliance issues; Mike Burke for his help in providing cumulative GPAs and with logistics needed to set up the study; Jenny Bergeron and Ellen Sarkisian of the Harvard University Derek Bok Center for Teaching and Learning for their assistance in collecting student survey information; and Annie Rota, Michael Hilborn, Arthur Barrett, and Jazahn Clevenger of the Harvard University Academic Technology Group for their assistance with the course learning management system.

