## Supplement for "Randomized controlled studies comparing traditional lectures versus online modules"

A neurological case was presented at Medical Grand Rounds. The patient, a 27-year-old man, had been diagnosed with the fragile X mental retardation syndrome. He had unusual facial features, including coarse acromegaloid features, large forehead, asymmetric long face, large ears, thick lips, and mandibular prognathism; impressive macroorchidism (enlarged testicles exceeding 100 mL volume); and hypotrophic (thin) lower extremities. His IQ was estimated to be less than 20. It was noted that this patient's presentation was unusually severe, much more dramatic than the typical case of fragile X syndrome (i.e., more severe macroorchidism, lower IQ, etc.).

The *FMR1* gene had been previously identified as the causal gene for fragile X syndrome, although in virtually all cases it was due to silencing of the *FMR1* gene as a result of a trinucleotide repeat expansion upstream of the gene. Gene sequencing of *FMR1* was performed for the patient. While there was no trinucleotide repeat expansion, there was a point mutation in the coding region of one allele of the gene, resulting in the alteration of isoleucine in position 367 to asparagine (Ile367Asn, or I367N).

1. What effect might the I367N mutation have on the protein's structure? How might this explain the unusually severe presentation of fragile X syndrome seen in this patient? Why might it be worse to have a point mutation of the gene rather than silencing of the gene?
   The mutation may cause the protein to unfold, since a hydrophobic residue is being replaced with a polar residue. It also might cause misfolding of the protein, or change the surface properties of the protein. The mutant protein may have adverse consequences that make it worse than having no protein; for example, the mutant protein may engage in inappropriate protein-protein interactions, or form aggregates that are toxic to cells. Generally speaking, the mutation may be worse than a loss-of-function—it may have some dominant-negative activity.

Computational algorithms were used to analyze the amino acid sequence of the FMR1 protein. Although it was not possible to determine the tertiary structure of the protein with computational analysis, the analysis did identify a distinct domain in the FMR1 protein surrounding the position of the mutation I367N. In collaboration with scientists, recombinant protein corresponding to this domain was produced in the laboratory. Recombinant protein with an I→N mutation in the appropriate position was also produced. Both the wild-type and mutant domains were subjected to circular dichroism (CD) spectroscopy:

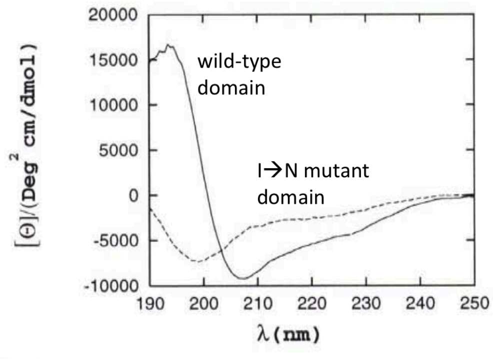


Adapted from Musco et al. (1997) *Nat Struct Biol* 4: 712-6

2. Based on this data, do you think the domain is predominantly made up of α-helices, β strands, or a combination of the two? What is the apparent effect of the I→N mutation on the structure of the domain?
   The curve of the wild-type domain is intermediate between the standard α-helix and β strand curves, suggesting the presence of both in the domain. The curve of the mutant domain is similar to that of random coil, suggesting that the mutant domain is entirely unfolded.

The scientists next attempted to determine the structure of the domain by X-ray crystallography. Despite much effort, they were unable to obtain high-quality crystals of the domain from FMR1. However, in searching through the genome, they found that there were a number of other proteins that appeared to have domains highly similar (homologous) to the FMR1 domain. Moreover, biochemical studies had shown that some of these other domains bound to RNA molecules. This suggested that FMR1 was an RNA-binding protein.

The scientists then worked with a homologous domain from a protein called Nova. They succeeded in obtaining crystals with this domain, acquiring X-ray diffraction data, and solving the three-dimensional structure (below). They also used homology modeling to speculate as to what the FMR1 domain might look like. The structure was notable for having a three-stranded β sheet on one "side" and three α-helices on the other "side."

Nova domain | Nova domain | FMR1 domain (homology model)

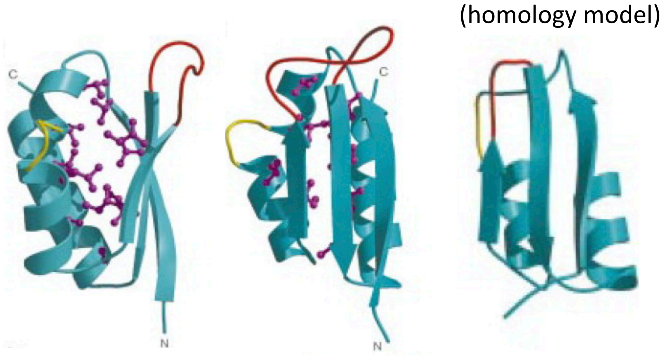


Hydrophobic side chains in the protein core are shown as ball-and-stick models. Adapted from Lewis et al. (1999) *Structure* 7, 191-203.

3. Based on this structure, where do you think the RNA binds to the protein? Along the β sheet? Across the α-helices? Somewhere else? (Take your best guess.)
   There is no right answer, as there is no way to know for sure. In real life, at the time, the scientists made bets on what the answer would turn out to be. They were all wrong.

With the crystal structure of the domain solved, the scientists then sought to understand how the domain interacts with RNA. They used a biochemical technique called SELEX to identify an RNA sequence that bound with high affinity to the domain. The resulting RNA sequence was 20 nucleotides in length, with the key portion of the sequence being bases 12-15, which were uracil-cytosine-adenine-cytosine (UCAC). They attempted to obtain crystals of the domain bound to this 20-nucleotide RNA.

In the interim, the scientists put together a sequence alignment of similar domains from Nova and related proteins from a variety of species (see next page). They used computational algorithms to help identify secondary structural elements and locations of gaps or insertions.

4. Looking at the alignment, which amino acids are identical in all of the domains? Which amino acids are similar (i.e., have conservative substitutions) in all of the domains? What does this imply about the importance of these amino acids? How many of these are hydrophobic amino acids that likely contribute to the hydrophobic core? Is there any particular non-hydrophobic amino acid that stands out?
   The only identical amino acids are Gly-22, Gly-25, and Arg-54. Other amino acids are well-conserved, e.g., position 33 is almost always a positively charged amino acid (Lys/Arg/His). Most of the well-conserved amino acids are hydrophobic and in the protein core. The most notable non-hydrophobic amino acid is Arg-54. Gly-22 and Gly-25 may be important for creating a turn between two α-helices.

| | | S1 H1 | I | H2 S2 | V | S3 H3 |
|---|---|---|---|---|---|---|
| | | 5...10...15...20. | ..25 | ...30...35...40.. | .45...50.. | .55...60...65...70...75. |
| | KH1 | QYFLKVLIPSYAAGSII | GKGG | QTIVQLQKETGATIKLS | KSKDFYPGTT | ERVCLIQGTVEALNAVHGFIAEKI |
| **Nova-1** | KH2 | ANQVKIIVPNSTAGLII | GKGG | ATVKAVMEQSGAWVQLS | QKPDGINLQ | ERVVTVSGEPEQNRKAVELIIQKI |
| | KH3 | KDVVEIAVPENLVGAIL | GKGG | KTLVEYQELTGARIQIS | KKGEFVPGTR | NRKVTITGTPAATQAAQYLITQRI |
| | KH1 | EYFLKVLIPSYAAGSII | GKGG | QTIVQLQKETGATIKLS | KSKDFYPGTT | ERVCLVQGTAEALNAVHSFIAEKV |
| **Nova-2** | KH2 | AKQAKLIVPNSTAGLII | GKGG | ATVKAVMEQSGAWVQLS | QKPEGINLQ | ERVVTVSGEPEQVHKAVSAIVQKV |
| | KH3 | KELVEIAVPENLVGAIL | GKGG | KTLVEYQELTGARIQIS | KKGEFLPGTR | NRRVTITGSPAATQAAQYLISQRV |
| **αCP-1/** | KH1 | TLTIRLLMHGKEVGSII | GKKG | ESVKRIREESGARINIS | EGNCP | ERIITLTGPTNAIFKAFAMIIDKL |
| **PCBP-1/** | KH2 | PVTLRLVVPATQCGSLI | GKGG | CKIKEIRESTGAQVQVA | GDMLPNST | ERAITIAGVPQSVTECVKQICLVM |
| **hnRNP E1** | KH3 | QTTHELTIPNNLIGCII | GRQG | ANINEIRQMSGAQIKIA | NPVEGSS | GRQVTITGSAASISLAQYLINARL |
| **αCP-2/** | KH1 | TLTIRLLMHGKEVGSII | GKKG | ESVKKMREESGARINIS | EGNCP | ERIITLAGPTNAIFKAFAMIIDKL |
| **PCBP-2/** | KH2 | PVTLRLVVPASQCGSLI | GKGG | CKIKEIRESTGAQVQVA | GDMLPNST | ERAITIAGIPQSIIECVKQICVVM |
| **hnRNP E2** | KH3 | TTSHELTIPNDLIGCII | GRQG | AKINEIRQMSGAQIKIA | NPVEGST | DRQVTITGSAASISLAQYLINVRL |
| | KH1 | TLTIRLLMHGKEVGSII | GKKG | ETVKKMREESGARINIS | EGNCP | ERIVTITGPTDAIFKAFAMIAYKF |
| **αCP-3** | KH2 | PVTLRLVVPASQCGSLI | GKGG | SKIKEIRESTGAQVQVA | GDMLPNST | ERAVTISGTPDAIIQCVKQICVVM |
| | KH3 | ASTHELTIPNDLIGCII | GRQG | TKINEIRQMSGAQIKIA | NATEGSS | ERQITITGTPANISLAQYLINARL |
| | KH1 | TLTLRMLMHGKEVGSII | GKKG | ETVKRIREQSSARITIS | EGSCP | ERITTITGSTAAVFHAVSMIAFKL |
| **αCP-4** | KH2 | PVTLRLVIPASQCGSLI | GKAG | TKIKEIRETTGAQVQVA | GDLLPNST | ERAVTVSGVPDAIILCVRQICAVI |
| | KH3 | TSSQEFLVPNDLIGCVI | GRQG | SKISEIRQMSGAHIKIG | NQAEGAG | ERHVTITGSPVSIALAQYLITACL |
| | KH1 | MVELRILLQSKNAGAVI | GKGG | KNIKALRTDYNASVSVP | DSSGP | ERILSISADIETIGEILKKIIPTL |
| **hnRNP K** | KH2 | DCELRLLIHQSLAGGII | GVKG | AKIKELRENTQTTIKLF | QECCPHST | DRVVLIGGKPDRVVECIKIILDLI |
| | KH3 | IITTQVTIPKDLAGSII | GKGG | QRIKQIRHESGASIKID | EPLEGSE | DRIITITGTQDQIQNAQYLLQNSV |
| | KH1 | TYHMKILVPAVASGAII | GKGG | ETIASLQKDTGARVKMS | KSHDFYPGTT | ERVCLITGSTEAIMVVMEFIMDKI |
| **Pasilla** | KH2 | DKQVKILVPNSTAGMII | GKGG | AFIKQIKEESGSYVQIS | QKPTDVSLQ | ERCITIIGDKENNKNACKMILSKI |
| | KH3 | KDSKNVEVPEVIIGAIL | GPSG | RSLVEIQHVSGANVQIS | KKGIFAPGTR | NRIVTITGQPSAIAKAQYLIEQKI |
| | KH1 | TLTIRLIMQGKEVGSII | GKKG | EIVNRFREESGAKINIS | DGSCP | ERIVTVSGTTNAIFSAFTLITKKF |
| **Mub** | KH2 | QIPIRLIVPASQCGSLI | GKSG | SKIKEIRQTTGCSIQVA | SEMLPNST | ERAVTLSGSAEQITQCIYQICLVM |
| | KH3 | QQQHEMTVSNDLIGCII | GKGG | TKIAEIRQISGAMIRIS | NCEEREGGNT | DRTITISGNPDSVALAQYLINMRI |
| | KH1 | EETVRILIPSSIAGAVI | GKGG | QHIQKMRTQYKATVSVD | DSQGP | ERTIQISADIESTLEIITEMLKYF |
| **Bancal** | KH2 | DFDVRLLIHQSLAGCVI | GKGG | QKIKEIRDRIGCRFLKVF | SNVAPQST | DRVVQTVGKQSQVIEAVREVITLT |
| | KH3 | NNSTQVTIPKELAGAII | GKGG | GRIRRIRNESSAYITID | EPLPNSN | DRIITISGTPKQIQMAQYLLQQSV |
| | KH1 | DVHLRMLCLVKHASLIV | GHKG | ATISRIKSETSARINIS | NNIRGVP | ERIVYVRGTCDDVAKAYGMIVRAL |
| **Pbp2p** | KH2 | EISINLLIPHHLMGCII | GKRG | SRLREIEDLSAAKLFAS | PNQLLLSN | DRILTINGVPDAIHIATFYISQTL |
| | KH3 | FVQQEIFIDEKFVGNVI | GKDG | KHINSVKESTGCSIIIQ | DPVEGSS | ERRLTIRGTFMASQAAIMLISNKI |

Notes: This is an alignment of putative RNA-binding domains from Nova proteins and other closely related proteins from a variety of species. Each of these proteins has a total of three RNA-binding domains, numbered 1 through 3. **The domain that was crystallized with RNA was the third domain of Nova-2.** The numbering at the top of the alignment thus follows the third domain of Nova-2. The last digit of each number is aligned with the amino acid in the position indicated by the number. Secondary structure predictions are shown at the top of the alignment. The arrows indicate β strands. The cylinders indicate α-helices. The letter "I" indicates a non-α, non-β region of invariant length (4 amino acids). The letter "V" indicates a non-α, non-β region of variable length.

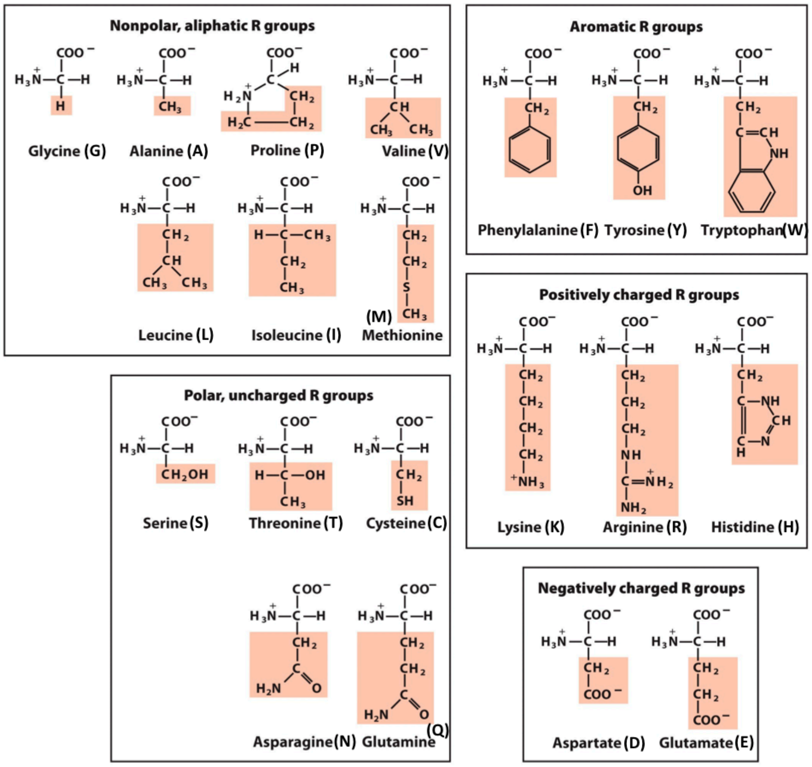

The scientists successfully determined the crystal structure of the Nova domain bound to the 20-nucleotide RNA. They discovered that the RNA actually binds along the “seam” of the protein, between the β sheet and α-helical side of the protein. The RNA bases, particularly the UCAC sequence, are positioned to make hydrogen bonds with amino acids in the protein.

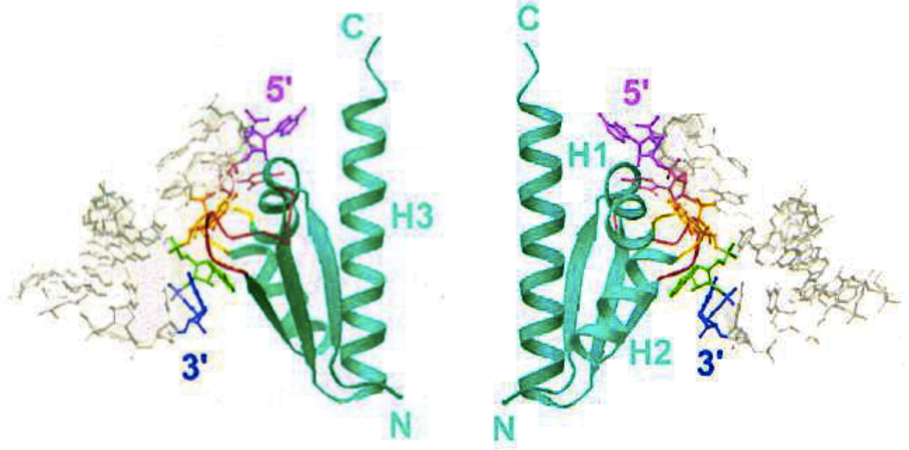


**Cyt-13 is the first C in UCAC**

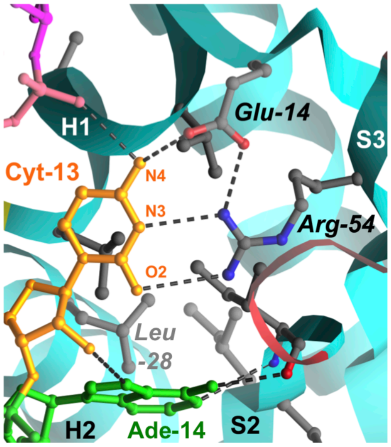


**Leu-28 is in the same position as the FMR1 I→N mutation**

5. How do Glu-14 and Arg-54 recognize the cytosine base (Cyt-13)? Do you think that Glu-14 and Arg-54 are critical for protein function? Looking at the alignment on the previous page, is there evidence that these amino acids are important?
   Glu-14 and Arg-54 coordinate to form three parallel hydrogen bonds with the cytosine, acting as a molecular mimic of guanine. This creates a pseudo-Watson-Crick basepair. Glu-14 and Arg-54 together appear to be essential for binding and recognizing a cytosine at that position in the RNA molecule. While Glu-14 is not particularly well conserved across domains, Arg-54 is found in every domain, suggesting that it is absolutely critical for the function of the proteins.

6. What is the most conservative possible amino acid substitution for arginine? What do you think would happen if you replaced Arg-54 with this amino acid? Would the protein still function properly?
   Lysine, which is also a long, positively charged amino acid. However, it is not as long as arginine, and whereas arginine has two nitrogen atoms at the end of the side chain, lysine only has one nitrogen atom. Thus, it is unlikely that lysine can form two or even one hydrogen bond with Cyt-13, and the domain would not bind well to the RNA molecule.

7. Leu-28 in the Nova domain is in the corresponding position to the FMR1 I→N mutation. Does it make sense for Nova to use a leucine, whereas FMR1 ordinarily uses an isoleucine in this position? Given the position of Leu-28 in the Nova domain with respect to the RNA, does it make sense that the I→N mutation in FMR1 should cause the FMR1 domain to unfold? What is an alternative explanation for how the I→N mutation causes FMR1 dysfunction and, thus, the patient’s disease?
   Leucine and isoleucine are structurally similar amino acids and so would represent a conservative substitution. Although at first glance it appears that Leu-28 is part of the hydrophobic core, it is actually exposed to the surface, and the RNA binding may actually be stabilized by the “stacking” of Cyt-13 and Ade-14 on the hydrophobic side chain. If changed to asparagine, it may actually not disrupt the hydrophobic core, but may instead simply impair RNA binding, thus causing a loss of protein function.

The scientists produced more of the Nova domain as well as a mutant version of the Nova domain with Arg-54 replaced with the conservative substitution (as asked in question 6). They performed CD spectroscopy of the wild-type and mutant Nova domains (left). They also assessed for binding of the domains to the 20-nucleotide RNA (right).

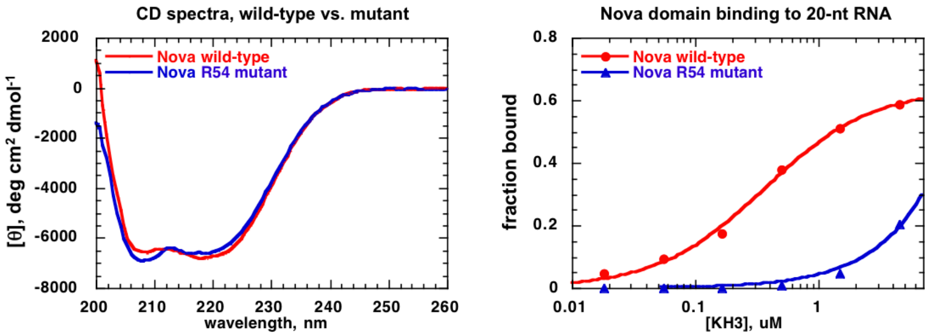


8. In light of these data, what is the effect of the conservative substitution on protein structure and function? Is this what you would have predicted from the crystal structure?
The mutant Nova domain is impaired in RNA binding, but this is not the result of unfolding of the domain. Instead, it appears to be purely the result of removing the hydrogen bonds ordinarily provided by Arg-54 to Cyt-13, as suggested by the crystal structure.

Given the competing models for how the I→N mutation might affect the structure and function of FMR1, the scientists decided to do a genetic experiment. They obtained *FMR1* null (knockout) mice, which are similar to human patients in that they have macroorchidism (enlarged testicles compared to wild-type mice). The scientists created a version of the *FMR1* gene encoding the I367N mutation and inserted it into *FMR1* null mice, thereby creating transgenic mice overexpressing the mutant *FMR1* gene (the mice ended up with several copies of the mutant *FMR1* gene incorporated into the genome).

The scientists took *FMR1* null mice and transgenic mice (*FMR1* null + *FMR1* I→N) that were littermates (so as to control for age, genetic background, and environmental conditions) and assessed their testicular size.

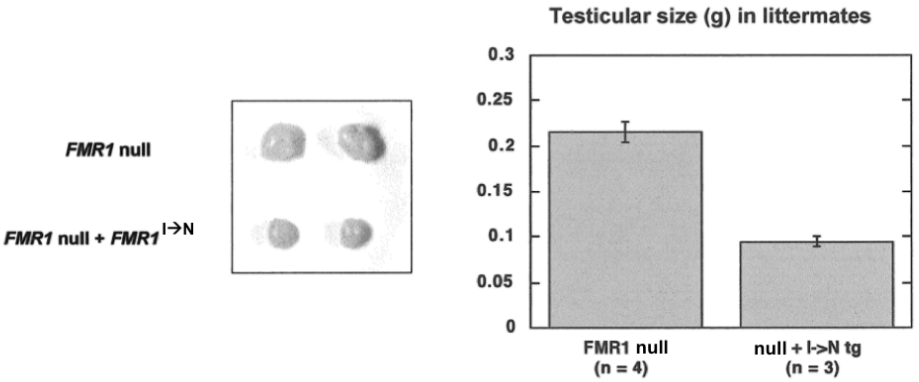


9. In light of these data, what is the effect of the I→N mutation on the function of the FMR1 protein? Is this consistent with the patient's severe presentation?
Mice that are overexpressing the mutant protein have smaller testicles. This implies that the mutation is actually a partial loss-of-function mutation, with multiple copies of the mutant gene resulting in reversal of the macroorchidism phenotype. If a full loss-of-function mutation (equivalent to the gene silencing normally responsible for fragile X syndrome), the mutant protein would not have changed the testicular size. If a dominant-negative mutation, the testicular size should have increased. Thus, the I→N mutation does not appear to be (solely) responsible for the patient's unusually severe presentation. Rather, there must be some other contributory factor(s) unique to this patient.
